# Re-weighting of Sound Localization Cues by Audiovisual Training

**DOI:** 10.1101/616490

**Authors:** Daniel P. Kumpik, Connor Campbell, Jan W.H. Schnupp, Andrew J King

## Abstract

Sound localization requires the integration in the brain of auditory spatial cues generated by interactions with the external ears, head and body. Perceptual learning studies have shown that the relative weighting of these cues can change in a context-dependent fashion if their relative reliability is altered. One factor that may influence this process is vision, which tends to dominate localization judgments when both modalities are present and induces a recalibration of auditory space if they become misaligned. It is not known, however, whether vision can alter the weighting of individual auditory localization cues. Using non-individualized head-related transfer functions, we measured changes in subjects’ sound localization biases and binaural localization cue weights after ~55 minutes of training on an audiovisual spatial oddball task. Four different configurations of spatial congruence between visual and auditory cues (interaural time differences (ITDs) and frequency-dependent interaural level differences (interaural level spectra, ILS) were used. When visual cues were spatially congruent with both auditory spatial cues, we observed an improvement in sound localization, as shown by a reduction in the variance of subjects’ localization biases, which was accompanied by an up-weighting of the more salient ILS cue. However, if the position of either one of the auditory cues was randomized during training, no overall improvement in sound localization occurred. Nevertheless, the spatial gain of whichever cue was matched with vision increased, with different effects observed on the gain for the randomized cue depending on whether ITDs or ILS were matched with vision. As a result, we observed a similar up-weighting in ILS when this cue alone was matched with vision, but no overall change in binaural cue weighting when ITDs corresponded to the visual cues and ILS were randomized. Consistently misaligning both cues with vision produced the ventriloquism aftereffect, i.e., a corresponding shift in auditory localization bias, without affecting the variability of the subjects’ sound localization judgments, and no overall change in binaural cue weighting. These data show that visual contextual information can invoke a reweighting of auditory localization cues, although concomitant improvements in sound localization are only likely to accompany training with fully congruent audiovisual information.

## INTRODUCTION

Accurate sound localization is achieved by integrating binaural cues (interaural level and time differences; ILDs and ITDs) and location-dependent spectral cues, which together constitute the head-related transfer function (HRTF). ITDs and ILDs provide information about a sound’s azimuth, whereas spectral cues are crucial for sound elevation judgements and for resolving front-back confusions (Wightman and Kistler, 1993; Jin et al., 2004). Depending on the frequency content of the sound and the egocentric location of its source, each of these cues alone may be spatially ambiguous, leading to localization errors (Blauert, 1997; Shinn-Cunningham et al., 2000; King et al., 2001). In line with work on Bayesian integration of both multisensory (e.g., Ernst and Banks, 2002; Alais and Burr, 2004; Körding et al., 2007; Rowland et al., 2007; Targher et al., 2012; Hollensteiner et al., 2015; Zhou et al, 2018) and unisensory (e.g., Landy and Kojima, 2001; Knill and Saunders, 2003; Hillis et al., 2004; Watt et al., 2005; Sturz and Bodily, 2010) spatial cues, the localizing brain can be usefully compared with an ideal observer, integrating sound localization cues to resolve ambiguities by weighting each cue according its relative reliability (Reijniers et al., 2014).

The relative weighting of individual auditory localization cues can change, as demonstrated by studies in which binaural cues are altered by temporarily occluding one ear (Kacelnik et al., 2006; Van Wanrooij and Van Opstal, 2007; Kumpik et al., 2010; Keating et al., 2013, 2016). These studies indicate that the auditory system can adapt rapidly to distortions in spatial hearing by giving greater weight to the cues that are less affected by the perturbation, i.e., the unchanged spectral cues provided by the non-occluded ear. Recent work in ferrets (Keating et al., 2013, 2015, 2016) and humans (Keating et al., 2016) has shown that such adaptation can be achieved either by up-weighting these cues or by learning a new relationship between the altered binaural cues and directions in space, depending on the spectral content of the stimuli used and therefore the localization cues that are available.

The importance of cue reliability for integration and remapping of sensory inputs has been demonstrated dramatically in multisensory studies that manipulate the dominance of vision over audition in spatial judgement tasks. For example, in the presence of a spatially displaced visual distractor, judgements of sound location are biased in the direction of the visual stimulus (the ventriloquism effect; Bertelson and Radeau, 1981). Similarly, using lenses to compress the visual field results in a corresponding change in the perception of auditory space (Zwiers et al., 2003). The ventriloquism illusion has been shown to reflect optimal integration of auditory and visual spatial cues, with the opposite effect – audition dominating audiovisual location judgements – occurring when visual information becomes spatially less reliable (Alais and Burr, 2004). Moreover, after exposure to spatially-discordant auditory and visual stimuli, subjects often show a shift of auditory localization in the direction of the previously presented visual stimulus (the ventriloquism aftereffect, VAE; Radeau and Bertelson, 1974; Recanzone, 1998). From a Bayesian perspective, this remapping of auditory space can be characterized as an updating of auditory likelihoods following exposure to spatially-conflicting but more reliable visual information (Wozny and Shams, 2011a).

Further evidence that cue reliability may determine how multisensory spatial information is integrated is provided by the demonstration that co-located visual cues can both improve the accuracy of sound localization judgments (Bolognini et al., 2007; Tabry et al., 2013; Hammond-Kenny et al., 2017) and contribute to the suppression of echoes (Bishop et al., 2011). Adaptive changes in sound localization accuracy following manipulation of auditory spatial cues can take place, however, in the absence of visual feedback (Kacelnik et al., 2006; Carlile and Blackman, 2014; Zonooz et al., 2019). Nevertheless, training paradigms that include visual cues can facilitate the ability of listeners to learn new associations between auditory spatial cues and directions in space (Strelnikov et al., 2011; Berger et al., 2018) and to utilize ILDs appropriately following bilateral cochlear implantation (Isaiah et al., 2014).

Together, these studies show that vision can have a profound impact on auditory localization, most likely due to its greater spatial reliability. Little is known, however, about the influence of visual inputs on behavioral sensitivity to different auditory spatial cues (Sarlat et al., 2006). Given that the relative perceptual weights of monaural and binaural cues can change following unilateral hearing loss, we sought to determine whether this is also the case when feedback about their relative reliability is provided by spatially congruent or conflicting visual signals. We performed four short-term training experiments that used different configurations of audiovisual spatial congruence to test the hypothesis that more reliable auditory localization cues undergo increases in perceptual weighting, whilst unreliable cues are down weighted. Our results provide insight into the changes in auditory processing that occur when auditory cues are spatially matched or mismatched with vision, and provide further support for the notion that acoustic cues for sound location are integrated in the brain in a statistically optimal fashion.

## MATERIALS AND METHODS

### Subjects

Nineteen subjects took part in the experiment (8 male, mean age ± SD: 22.7 ± 2.6 years). All had normal or corrected-to-normal vision and normal audiometric thresholds from 125 Hz to 8 kHz. The four audiovisual experiments were run in random order and were completed on different days. Subjects were recruited through online and departmental notices; all received payment for their time and provided informed consent before beginning the study. Ethical approval was provided by the Medical Sciences Inter-Divisional Research Ethics Committee of the University of Oxford (study R52936).

### Apparatus

Subjects sat on a stool in a sound-attenuated chamber and a chin-rest was used to keep their head stationary during the experiment. Virtual auditory space stimuli were passed to a MOTU 828 MKII audio interface and presented to subjects over Sennheiser HD650 headphones. A Viewsonic PJD5453S projector, with a maximum resolution of 1920×1080 pixels, was mounted on the wall behind and above the subject and projected stimuli onto a semicircular screen that curved around the subject and covered azimuths up to 70° to the left and right of the midline. The screen was positioned at a radius of 84 cm from the center of the subjects’ heads, and extended from 88 cm-144 cm above the floor (thus covering 36.9° of visual field from top to bottom). It was composed of black speaker cloth mounted on a wooden frame, and a black cotton curtain was velcroed to the bottom of the frame to hide the metal legs. During the experiments, the azimuthal location of the visual and auditory stimuli was varied as described in further detail in the following section, while the vertical location (elevation) was held fixed at 116 cm above the floor, corresponding roughly to eye level.

### Stimuli

#### Auditory stimuli

Broadband Gaussian noise pulses were generated on each trial and were bandpass filtered and convolved with appropriate spatial cues according to the type of psychophysics run being conducted (see below). All auditory stimuli were presented over headphones calibrated to an RMS average binaural level of 70 dB SPL, using a Brüel and Kjær 4191 condenser microphone placed inside a Brüel and Kjær 4153 artificial ear and connected to a Brüel and Kjær 3110-003 measuring amplifier. Sinusoidal amplitude modulation (SAM) was applied at a rate of 6 Hz; in the sound localization and cue trading tasks, stimuli spanned 8 cycles of the modulator and were therefore 1.33 s in duration. In the spatial oddball task used for audiovisual training, between 6 and 12 noise stimuli were presented per trial with an inter-stimulus interval of 200 ms; each noise pulse was a single cycle of the modulator and therefore had a duration of 166.7 ms.

To produce realistically externalized auditory stimuli and thus encourage multisensory integration, we selected non-individualized HRTFs from the SYMARE database (Jin et al., 2014), and applied reverberations using a custom-designed room simulator, which added specular reflections of up to third-order to the presented sounds. The SYMARE database comprises head-related impulse response filter measurements for 393 source directions, sampling auditory space in ≤10° intervals from 45° below the horizon. We psychophysically determined which of the 10 individual HRTFs available in the database was best for each subject before training them on a sound localization task with the chosen HRTF (see *Procedure*). The ITDs for each virtual direction were computed using cross-correlation, and then adjusted as required for each experimental condition. We were thus able to manipulate the frequency-dependent ILDs, which incorporate the spectral cues at each ear (and following MacPherson and Middlebrooks (2002) are referred to here as interaural level spectra, ILS) independently from the ITD cue.

We combined sound localization cues in three general configurations that were used for different phases of the study. For HRTF selection, ILS-only localization training and ITD-only localization training, we presented ILS and ITD cues in isolation. The ILS-only localization stimuli were used in the first (familiarization) session to select an appropriate HRTF for each subject, and then to train subjects with the ILS cues from that HRTF. They were also used to re-familiarize subjects with the ILS cues before each audiovisual training experiment (see *Procedure*). ITD-only stimuli were only used for localization training purposes. ITDs are spatially ambiguous for high-frequency stimuli, whereas free-field ILDs (and spectral cues) tend to be negligible for low-frequency stimuli, a dichotomy known as the Duplex theory of sound localization (Strutt, 1907). The ILS stimuli were therefore band-pass filtered from 1.9 kHz-16 kHz and the ITD stimuli were band-passed from 0.5 kHz-1.3 kHz, in both cases to avoid frequency regions in which the other cue (which had a value corresponding to 0°) could play a prominent role in localization. To ensure that training was confined to the basic localization cues, reverberation was not added to these stimuli.

The second stimulus configuration used ILS cues and ITDs that were always congruent, while the third stimulus configuration employed ILS cues and ITD cues that were in spatial conflict. In both cases, broadband noise (0.5 kHz-16 kHz) was used as the stimulus and reverberation was applied.

#### Visual stimuli

All visual stimuli and markers were white on a black background. Geometric correction was applied to take account of the curvature of the screen. We used three types of visual stimuli: 1) a 0.48° wide fixation dot, which was located in front of and above the subject at 0° azimuth and 13.4° elevation; 2) a 1.23° wide cross hair mouse cursor, which subjects could control to initiate trials by clicking on the fixation dot, or respond on each trial by clicking on the location associated with the perceived sound-source direction; 3) during both pre-experimental sound localization training and experimental audiovisual training runs, we presented visual stimuli at specific angles relative to the horizontal position of the sound source in the form of blobs with a Gaussian contrast profile that covered 0.61° of visual field at full width half-maximum. During auditory-only localization training, the feedback stimuli were presented after the subjects’ response for a period of 400 ms, while during experimental audiovisual runs, visual stimuli were presented concurrently with the auditory stimulus throughout the 1.33 s duration of the sound and ramped up and down using a temporal contrast envelope that was aligned with the auditory amplitude modulation. Visual feedback stimuli were always presented at 0° elevation.

### Psychophysical tasks

Psychophysical tasks were controlled using Matlab (r.2015b; Mathworks) and its associated Psychophysics Toolbox (Brainard, 1997; Pelli, 1997; Kleiner et al., 2007).

#### Sound localization task

This task (Figure 1A) was used during HRTF selection, ILS and ITD training, and during the pre- and post-audiovisual training test session, with the session type determining the sound localization cues included in the stimuli and whether or not visual error feedback was provided (see *Procedure*). Subjects were seated in the chamber at the centre of the semicircular screen and fitted with headphones. They initiated trials by clicking the cross hair mouse cursor on the fixation dot. After 200 ms, the fixation dot disappeared and an externalized auditory stimulus was presented. Subjects were instructed to direct their gaze in the perceived direction of the sound source, and then respond by moving the cross hair cursor to that location and clicking the mouse button. Once the sound had ended, the fixation dot reappeared in preparation for the next trial.

**Figure 1.**
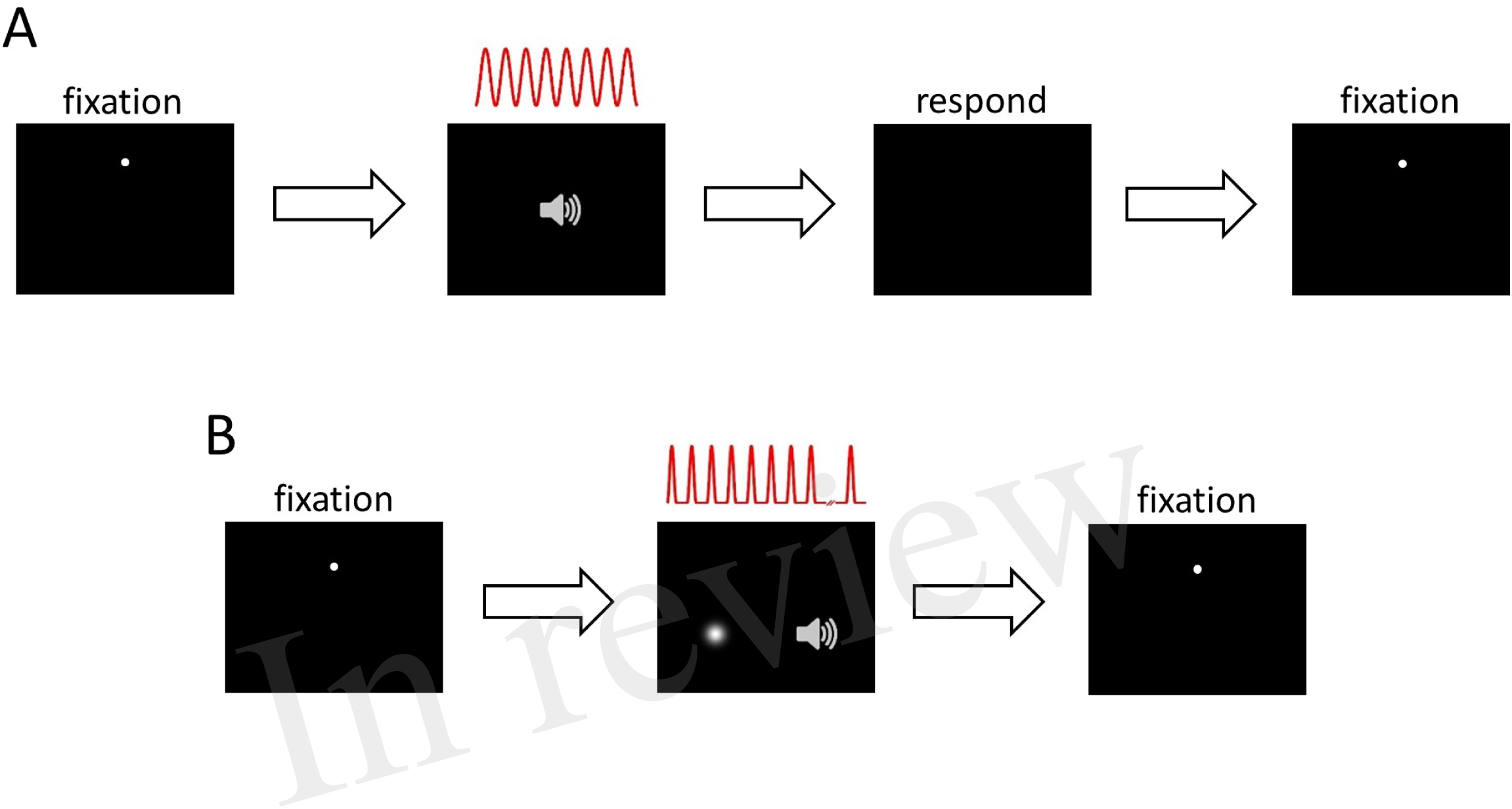
Psychophysical task structure. **(A)** Sound localization. Panel 1, inter-trial interval: subject fixates on white dot, moves mouse cursor over it and left-clicks to initiate trial. Panel 2, stimulus presentation: 200 ms after trial onset, 8 cycles of a sinusoidally amplitude modulated (SAM) auditory noise stimulus were presented, depicted in red. The auditory stimulus is spatialized by introducing ILS, ITDs or both cues. Panel 3, response: after stimulus presentation, subject makes an eye movement in the perceived direction of the auditory stimulus, then moves the mouse cursor over that position and left-clicks to respond. Panel 4, inter-trial interval and start of next trial. **(B)** Audiovisual spatial training. Panel 1, as in (A). Panel 2, stimulus presentation and spatial oddball detection: 200 ms after trial onset, a sequence of up to 12 spatialized auditory SAM cycles, separated by 200 ms, was presented concurrently with a visual “teacher” signal that indicated the reliability of the auditory spatial cues (auditory amplitude and visual contrast profile shown in red; see Methods). Subjects were required to ignore the auditory stimuli, monitor the visual stimulus for a spatial “oddball” that occured between the 6^th^ and the 11^th^ (inclusive) stimulus in the sequence, and to left-click when they detected the oddball. At this point stimulus presentation was halted. Panel 3, inter-trial interval and start of next trial.

#### Audiovisual training task

We used an audiovisual oddball detection task (Figure 1B) to manipulate the perceptual reliability of auditory localization cues, with the position of the visual stimulus acting as a “teacher” signal for recalibration of auditory space. Trials were initiated in the same way as the localization task. 200 ms after the fixation dot and mouse cursor had disappeared, a sequence of up to 12 auditory stimuli was presented, each consisting of a single modulation cycle (166.7 ms duration and separated by 200 ms) of the SAM noise stimulus. Concurrently with each auditory stimulus, a visual “Gaussian blob” stimulus (i.e., a bright patch with brightness fading at the edges according to a 2-D Gaussian spatial luminance profile) was presented, with the blob’s contrast rising and falling in synchrony with the amplitude modulation of the auditory stimulus. Eleven of the visual stimuli were presented at a single standard location in each trial (−20°, −10°, 0°, 10° or 20° azimuth; negative angles represent stimuli to the left of the midline), but one was a spatial oddball that could randomly appear as the 6^th^ to the 11^th^ stimulus in the sequence. The oddball was offset by either 10° or 20° to the left or right of the standard stimuli. Subjects were asked to direct their gaze to the standard visual stimulus sequence and then respond by clicking the mouse button when they detected the oddball. On each audiovisual presentation, the spatial relationship between the ILS and ITD cues and the visual stimuli was manipulated in different ways, depending on the training experiment (see *Procedure*). Each of the possible combinations for the standard and oddball angles was presented once, giving 20 trials per audiovisual training block. There were 20 blocks during the audiovisual training phase of each experiment, with subjects receiving a 30 s break between blocks. Subjects remained in the chamber with the door closed for the duration of the audiovisual training phase (~50 minutes) and subsequent post-training localization test (~5 minutes), and were asked to keep the headphones on during this period. The mean number of individual stimulus presentations per audiovisual training session was 8.5 ± 0.4 across all subjects and conditions.

### Procedure

#### HRTF selection phase

The first two-hour session of the experiment involved two phases, HRTF selection and auditory-only localization training. It was first necessary to determine, for each subject, which HRTF produced the most veridical sound localization performance. Subjects therefore completed:

i) A single ILS-only localization run with ITDs fixed at 0 µs, using a randomly chosen HRTF and high-pass noise (1.9 kHz-16 kHz), to familiarize them with the setup and the localization task. ILS cue values corresponding to stimulus angles of −20°, −10°, 0°, 10° and 20° (15 repetitions/angle) were used and visual feedback to stimulus location was given after each trial.

ii) HRTF selection runs with each HRTF tested in random order, ITDs fixed at 0 µs. Stimulus angles were −20°, −10°, 0°, 10° and 20° (10 repetitions/angle) and no visual feedback was provided. Sound localization performance on this task identified which HRTF provided the most veridical ILS cues, as determined by the HRTF that produced a slope closest to 1°/° when response angle was regressed on simulated stimulus angle.

#### Auditory localization training phase

Once the most appropriate HRTF had been selected, the first session concluded with a sound localization training phase, designed to familiarize subjects with the chosen non-individualized ILS cues. They also performed an ITD localization task during this phase to provide exposure to the range of ITDs presented over headphones. Subjects completed:

i) ILS-only training with visual feedback after each trial, using high-pass noise (1.9 kHz-16 kHz). Stimulus angles of −20°, −10°, 0°, 10° and 20° were used (15 repetitions/angle).

ii) Familiarization with ITD-only localization using band-pass noise (0.5 kHz-1.3 kHz). We used ITDs up to 145.8 µs on either side of the midline in 21.8 µs steps (5 repetitions/ITD). Because of the consistent relationship between ITDs and frontal sound locations, no visual feedback was provided in these runs, but subjects were informed that the stimuli could now come from anywhere within this range. The task was run repeatedly to reduce the mean square error (MSE) of the fit when response angle was regressed on ITD.

#### Test phase - matching ITDs to simulated HRTF angles

In the first hour of each of the following two-hour experimental audiovisual training sessions, subjects first performed localization tasks that were intended to re-familiarize them with the auditory stimuli and also to determine a set of ITDs to match the ILS cue angles to be used in the experimental audiovisual training block. This first hour involved:

i) Several ILS-only localization runs using high-pass noise (1.9 kHz-16 kHz) and stimulus angles of −20°, −10°, 0°, 10° and 20° (15 repetitions/angle). Visual feedback was provided after each trial and this task was intended to re-familiarize subjects with the non-individualized ILS cues.

ii) A single ILS-only localization run using high-pass noise (1.9 kHz-16 kHz) and stimulus angles of −20°, −10°, 0°, 10° and 20° (10 repetitions/angle). Visual feedback was not provided (Figure 2A).

iii) Several ITD-only localization runs with band-pass noise (0.5 kHz-1.3 kHz). We used ITDs up to ±145.8 µs in 21.8 µs steps (5 repetitions/ITD), and no visual feedback. This task was used to obtain ITDs that most closely matched subjects’ perceived angles of −20°, −10°, 0°, 10° and 20° (Figure 2B).

**Figure 2.**
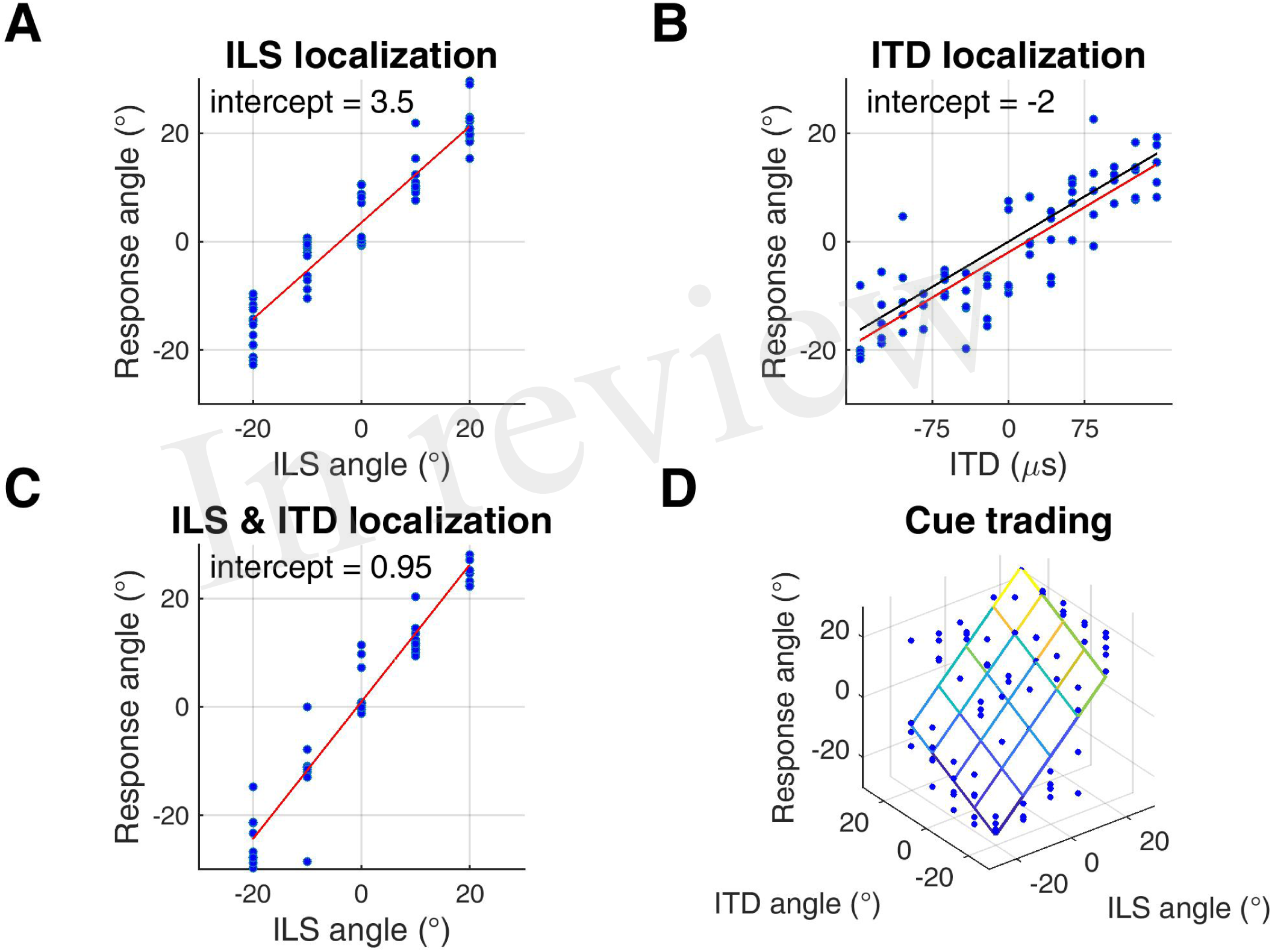
Sound localization tasks. **(A-C)** sound localization with ILS cues only (A), ITDs only (B), and ILS cues combined with spatially congruent ITD cues (C). Red lines are fits to the data and intercepts in top left indicate subjects’ response biases. In (B), the black line represents the centered function used to match ITDs to angles (see Methods). **(D)** Sound localization in the auditory spatial cue trading task carried out before and after audiovisual training. The fit illustrates a two-factor multiple regression of response angle on ILS angle and ITD angle.

Once a subject’s performance had stabilized in these tasks, as indicated by good and reproducible linear fits to the data across runs, we calculated the slope of a linear fit to the ITD localization run with the lowest MSE from the data collected that day. We used this ITD localization slope to calculate ITDs that best matched perceived angles of −20°, −10°, 0°, 10° and 20°; before doing so we removed the intercept of the fit to avoid introducing systematic biases into the ITD values (Figure 2B). After ILS and ITD training on the day of a subjects’ first audiovisual training experiment, we checked that sound localization performance with these combined cue values was reasonably veridical by having subjects perform a localization run with 10 repeats of each of the congruent ILS and ITD values. This run employed broadband noise (0.5 kHz-16 kHz) with reverberation added (Figure 2C). These stimuli therefore provided naturalistic virtual acoustic space (VAS) cues, and were used for all stimuli from here onwards.

Subjects’ ILS-only and ITD-only training performance in subsequent experimental sessions tended to stabilize quickly. Therefore, to mitigate subject fatigue before an experimental audiovisual training block, we did not reassess combined-cue localization in subsequent experiments unless the single-cue training data appeared anomalous due to, for example, the subject attending the session with a cold. On these rare occasions retraining was performed and/or the audiovisual training experiment was abandoned and re-run at a later date.

#### Experimental phase – audiovisual spatial training

In the second hour of each test session, subjects first performed a pre-training auditory localization test (ILS vs. ITD cue trading). This was followed by an audiovisual spatial training task (spatial oddball detection), and the session concluded with a post-training auditory localization test (ILS vs. ITD cue trading). The tasks were as follows:

#### Cue trading (pre/post-audiovisual training)

This involved measuring sound localization with stimuli containing ILS and ITDs, and with every combination of the 5 test angles for the two cues. There were therefore 25 different stimuli and each was repeated 5 times. This allowed us to determine spatial gains for the individual cues (i.e., how each cue mapped to spatial location, given the presence of the other cue) using a two-factor multiple regression of response angle on ILS angle and ITD angle (see Figure 2D). The regression intercept corresponded to a subject’s lateral localization bias, i.e., the distance of their mean responses from 0°. This task was performed before and after the audiovisual spatial training task and subjects were instructed that the stimuli could come from anywhere within the horizontal angular range of the stimuli (i.e., ±20°).

#### Audiovisual spatial training (spatial oddball detection) tasks

*i) Audiovisual congruent (perceptual training; AVCON)*. The ILS and the ITDs were always congruent with the visual stimulus.

*ii) ILS training with unreliable ITD (random-ITD)*. The ILS cue was always congruent with the visual stimulus. The ITD cue was presented from a position up to 20° off the midline that was selected from a uniform random distribution at 10° intervals and changed for each stimulus within a trial.

*iii) ITD training with unreliable ILS (random-ILS)*. The ITD was always congruent with the visual stimulus. The ILS cue was presented from a position up to 20° off the midline that was selected from a uniform random distribution at 10° intervals and changed for each stimulus within a trial.

*iv) Audiovisual remapping (ventriloquism aftereffect; VAE)*. The ILS and ITD cues were always congruent with each other, but the auditory stimulus was offset to the left or right of the visual stimulus by 10° for each stimulus within a trial; the direction of offset was decided at random for each subject, although approximately equal numbers were trained on each side of space.

### Data analysis

To check whether the use of non-individualized HRTFs produced cue trading responses that were not well-described by a simple two-cue linear model, we assessed the fit of the model to the pre-audiovisual training data for each subject. We pooled subjects’ data across their pre-test sessions and fitted the linear regression model given in Equation 1. We then used the model coefficients to predict subjects’ localization responses for the unique ILS and ITD cue conflict angle combinations used in the experimental phases of this study. The predictions from the model, and the mean angular response for each subject, are shown in Figure 3. For most subjects, a simple linear cue integration model is a reasonable description of their sound localization performance. However, the model performs poorly for several subjects, possibly caused by systematic cue biases that arose when one binaural cue had an intrinsically substantially higher spatial gain (and was therefore weighted more) than the other (see Figure 4).

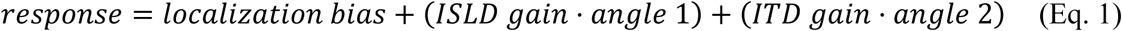

**Figure 3.**
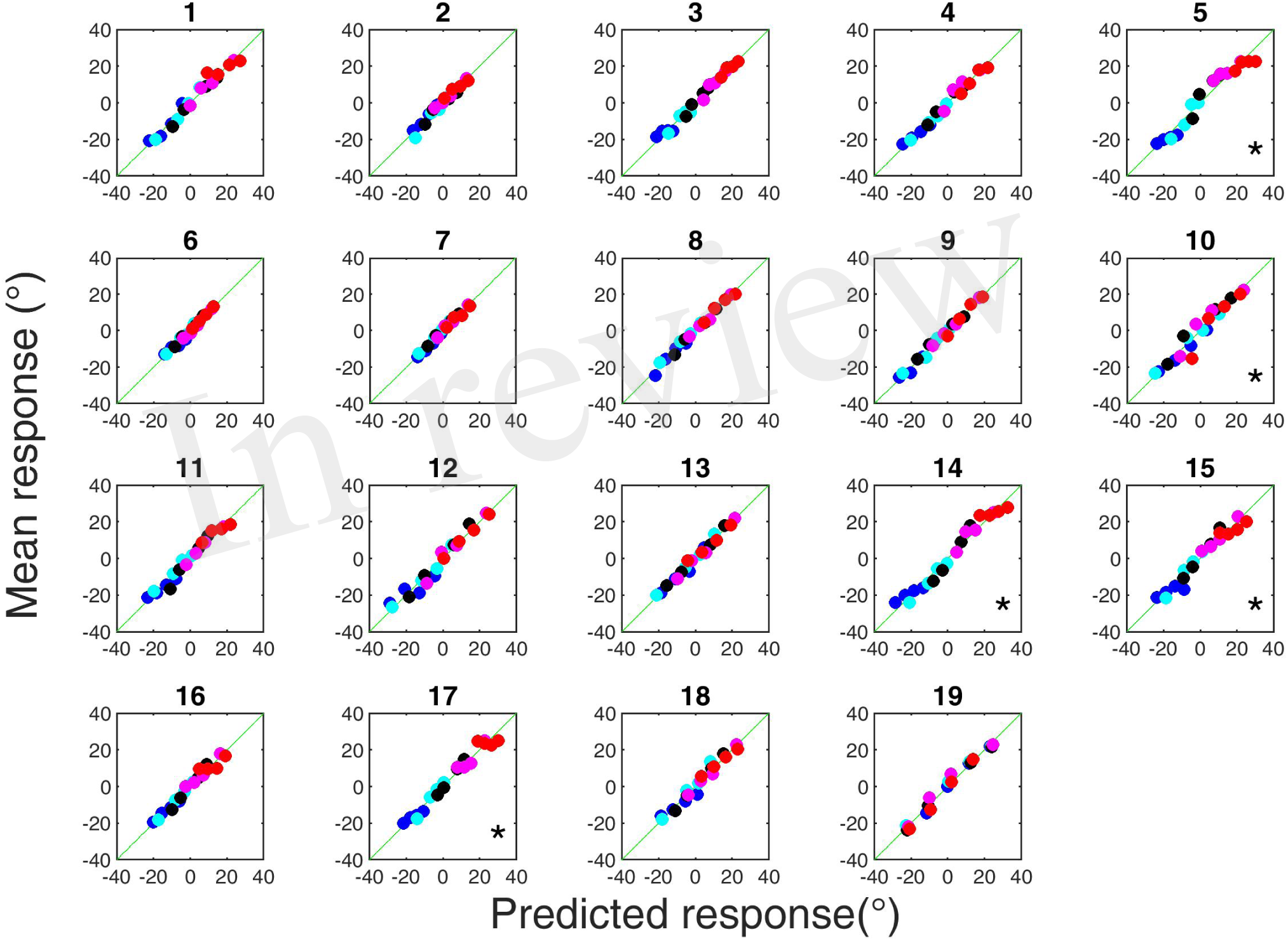
Subjects’ mean pooled pre-training responses to conflicting ILS and ITD cue values as a function of responses predicted from the same datasets using the linear cue integration model. Asterisks in the bottom right corner indicate datasets that were not considered for further analysis according to our exclusion criteria (see Methods). Symbols of different colors indicate different ILS positions; symbols of the same color are the different ITD positions that were presented at that ILS.

**Figure 4.**
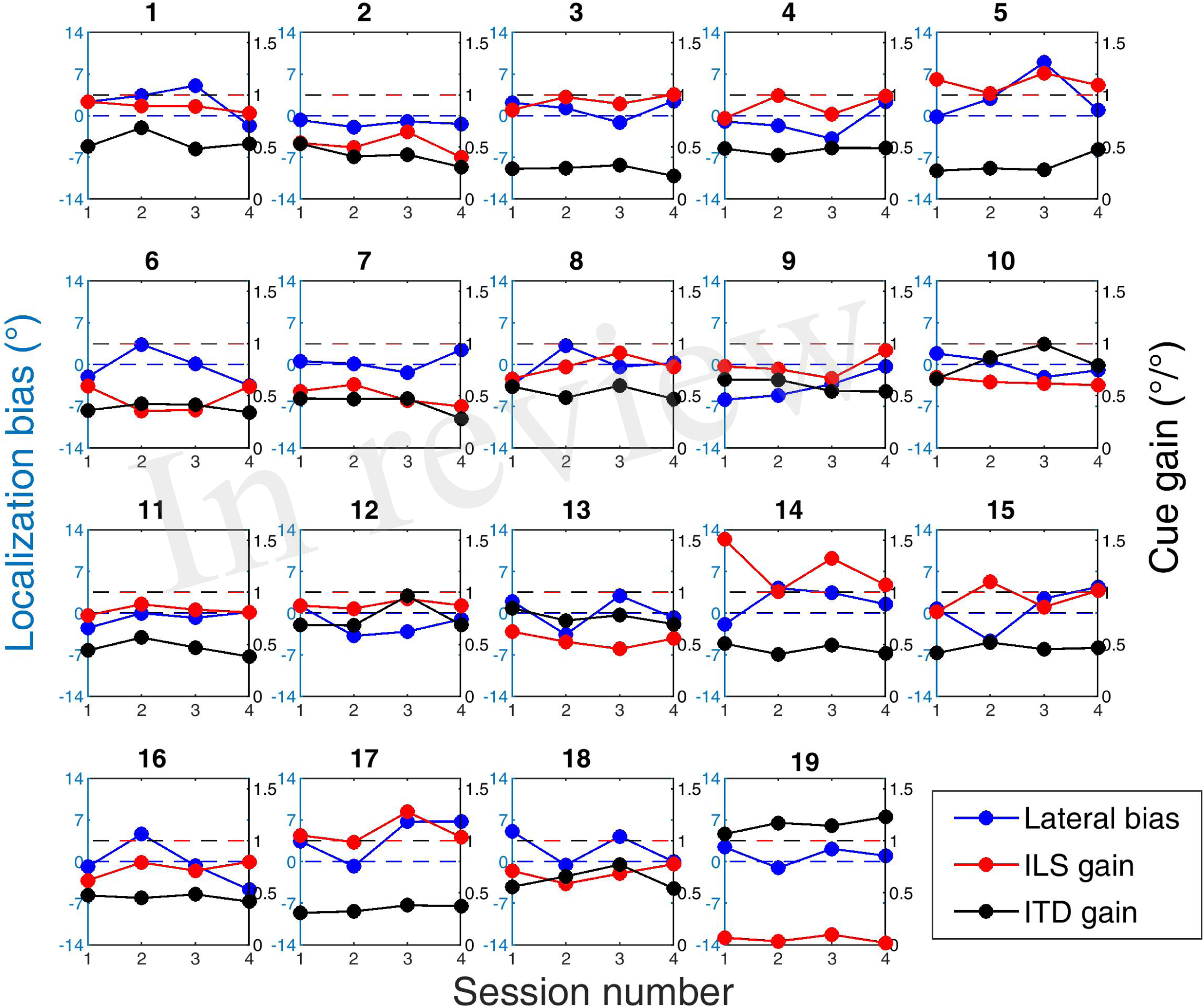
Subjects’ pre-training cue trading regression parameters across experimental sessions. Blue, localization bias; red and black, ILS and ITD cue gains, respectively. The horizontal blue dashed line indicates a localization bias of 0° and the red and black dashed lines denote an ILS or ITD gain of 1, respectively.

In order to assess the significance of these model violations, we fitted a regression to the mean predicted scores and checked each subject’s residuals for normality using a Kolmogorov-Smirnov single-sample goodness-of-fit test. This test showed that the linear cue integration regression model fitted 14 of our 19 subjects well, but the residuals for subjects 5, 10, 14, 15 and 17 deviated significantly from normal, indicating that the model was a poor fit to their pre-training cue trading data (Figure 3). We therefore excluded the data from those subjects from further analysis.

For each audiovisual training experiment, we obtained regression intercepts and ILS and ITD slopes from individual pre- and post-training cue-trading test blocks by performing a robust regression using the model given in equation 1. We determined significant group differences between spatial ILS and ITD cue gains before training by fitting a linear mixed-effects (LME) multiple regression model to the pre-training cue trading data, with ILS and ITD angle as fixed predictors (as in Equation 1) and with a random effect for the intercept, ILS slope and ITD slope for each subject. We analyzed group parameter changes within each experiment using a similar LME model that also included an interaction term for test session (pre-training vs. post-training). Within-subject training-induced changes were assessed by fitting each subject’s localization data to the regression model given in Equation 1, with an additional interaction term for test session (pre-training vs. post-training) included. We then performed ANOVA on the model coefficients.

From here on, we refer to the regression intercept as “localization bias”, and to the regression slopes as “spatial cue gains”. We identified general trends in bias and gain changes by regressing post-training biases and gains on those obtained before audiovisual training after removing outliers using a leverage test (Hoaglin and Welsch, 1978). To characterize underlying within-subject changes in ILS and ITD spatial gains, we calculated the change in ITD spatial gains as a function of the respective change in ILS spatial gain. We then bootstrapped the origin of a 95% confidence ellipse around the data by resampling the paired dataset with replacement 10,000 times, and calculated maxima for the joint distribution of changes in ILS and ITD spatial gains. To determine how spatial gain changes translate into the relative weighting of ILS and ITDs, we derived a “binaural weighting” index by expressing subjects’ pre- and post-training ILS spatial gains as a proportion of the summed gains from both cues in each pre- or post-training cue trading run and subtracting 0.5 from the resulting coefficients. According to this definition, binaural weighting values of −0.5 and 0.5 represent the extreme situations where either ITD or ILS dominate, respectively, and a value of 0 represents equal weighting between the cues. The threshold for determining statistical significance in all analyses was set at *p* < 0.05.

## RESULTS

### Training with spatially congruent auditory and visual cues (AVCON)

In this experiment, subjects performed the audiovisual spatial oddball task with spatially congruent sound localization cues and visual stimuli. Given that brief exposure to audiovisual stimuli can improve auditory localization with non-individualized HRTFs (Berger et al., 2018), we hypothesized that the presence of the visual stimuli would lead to an improvement in sound localization in the form of a reduction in the magnitude and variability of subjects’ localization biases. We also hypothesized that this would increase the spatial gain of (i.e., reduce response undershoot in) the more reliable binaural cue, with no change or a reduction in the spatial gain of the less reliable cue.

Eleven subjects showed reductions in localization bias (seven significant; Figure 5A) and three increased their biases (two significant). The slope of the regression of post-on pre-training localization bias shown in Figure 5A is not significantly different from 0 (0.2, t = 1.82, *p* = 0.11), and the magnitude of the change was negatively correlated with subjects’ pre-training biases (Pearson’s r correlation coefficient = −0.94, *p* < 0.001) (see Figure 9A). In other words, the greater the initial bias, the larger the bias reduction that occurred during spatially congruent audiovisual training. Overall, the group mean localization bias shifted leftwards (pre-training mean bias ± SD, −0.07 ± 2.84°; post-training, −0.95 ± 1.63°), a significant net change (linear mixed effects (LME) model on group data: t(3494) = −4, *p* < 0.0001). This effect is less apparent when high-leverage outliers are disregarded, as indicated by the non-significant intercept of the regression of post-training on pre-training localization bias (−0.49°: 95% confidence bounds −1.03, 0.05, t = −2.09, *p* = 0.07; Figure 5A). Thus, although training with spatially congruent visual and auditory cues reduced subjects’ localization biases in most cases, our data contain hints that their mean performance converged to a point just to the left of the midline, rather than at the midline itself.

**Figure 5.**
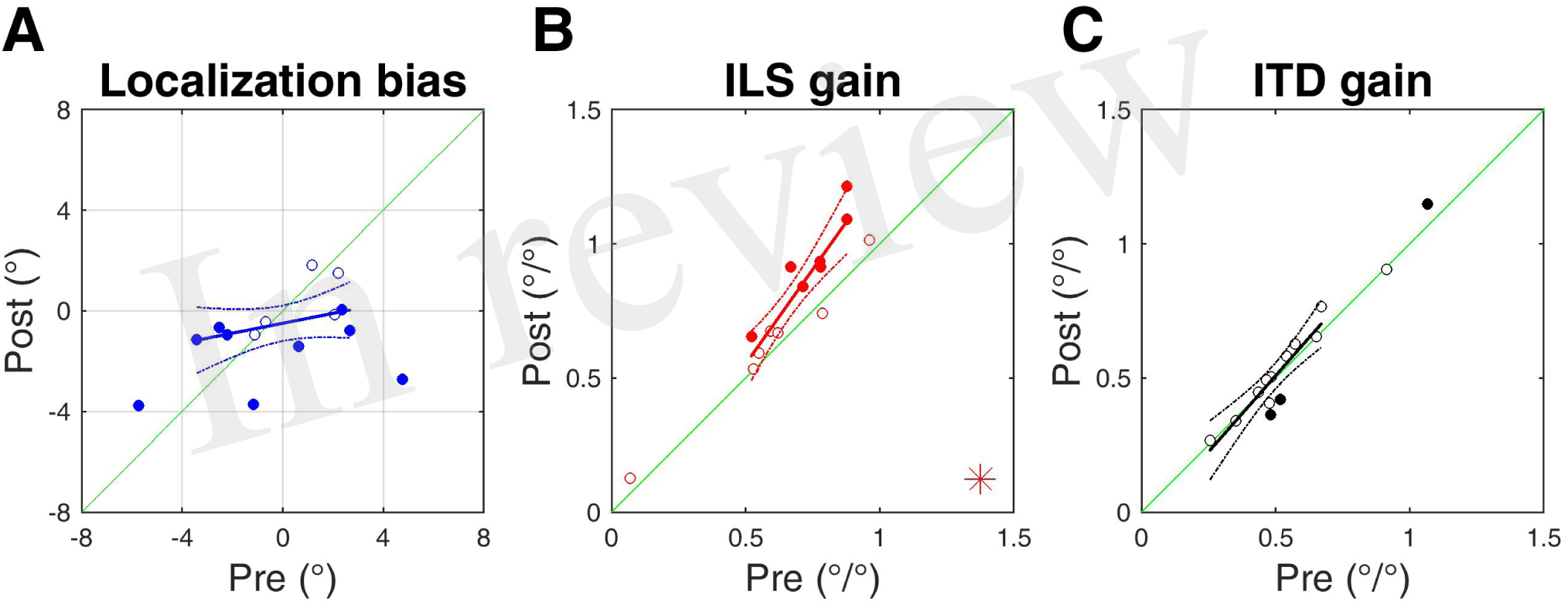
Subjects’ post-training multiple regression coefficients as a function of their pre-training regression coefficients for the AVCON experiment. **(A)** Localization bias. **(B)** ILS cue gains. **(C)** ITD cue gains. Asterisks in the bottom right corner indicate significant group effects of training according to a linear mixed effects model. Solid symbols indicate significant within-subject changes (see Results). Solid lines are regressions of post-training score on pre-training score after outliers were excluded using a leverage test. Dashed lines are 95% confidence intervals for the fits.

We next examined the post-training ILS and ITD spatial gains as a function of pre-training gain (Figure 5B, C). Prior to training, the spatial gains for all but two subjects were higher for ILS than ITDs (mean ILS gain ± SD, 0.67 ± 0.22°/°; ITD, 0.56 ± 0.21°/°). Since the very high ITD values from those two subjects caused them to be outliers according to a leverage test, we fitted an LME model to the pre-test response data and found this difference to be significant (adjusted LME estimate [and 95% confidence bounds]: ILS, 0.72°/° [0.61, 0.84]; ITD, 0.56°/° [0.45, 0.67]). Following audiovisual training, thirteen out of fourteen subjects increased their ILS gains (Figure 5B), and in seven individuals this increase was significant (the change for one additional subject approached significance (*p* = 0.055). Of those eight subjects whose ILS gain increases were significant or near-significant, seven also exhibited significant reductions in their localization biases. There was, however, no correlation between the change in localization bias and the change in ILS gain (r = −0.08, *p* = 0.79). The mean ILS gain increased from 0.67 ± 0.22°/° to 0.78 ± 0.27°/°, a group change that was significant (LME model with interaction term for pre-test vs. post-test; t(3494) = 7.38, *p* < 0.0001). The slope of the fit shown in Figure 5B is significantly greater than 1 (1.41 [1.02, 1.8]; t = 8.4, *p* < 0.0001), indicating that the largest increases in ILS gain occurred in subjects for whom this cue was already a reliable indicator of sound location. Exposure to spatially-congruent audiovisual cues resulted in a significant decrease in ITD gain in two subjects, an increase in one, and non-significant changes in the other nine (Figure. 5C); as a group there was no change (pre-training gains, 0.56 ± 0.21°/°; post-training, 0.57 ± 0.24°/°; t(3494) = −0.007, *p* = 0.65). Taken together, these data suggest that the weighting of ILS, presumably the more reliable sound localization cue, was increased relative to ITDs following repeated exposure to spatially-congruent auditory and visual information.

### Audiovisual training with congruent visual and ILS cues and randomized ITDs (random-ITD)

For this experiment, we manipulated the relative reliability of ILS and ITD localization cues by spatially pairing the ILS cue with the visual stimuli, whilst the ITD value was selected at random on each stimulus presentation. We therefore expected ILS spatial gains to increase whilst a decrease in gain was predicted for ITDs, resulting in a net increase in ILS weight. We hypothesized that these changes would be accompanied by a reduction in localization bias as subjects underwent reliability-driven perceptual learning for localizing the more dominant ILS.

Two subjects underwent significant decreases in their auditory localization biases and two subjects showed significant increases (Figure 6A), whereas there was essentially no change in the others. As a group, there was a significant leftwards shift in bias (pre-training mean bias, −0.02 ± 2.66°; post-training, −0.76 ± 3.03°; LME model: t(3494) = −3.25, *p* < 0.01). However, removal of three high-leverage outliers revealed that audiovisual training on this task did not change localization accuracy: subjects generally retained their initial biases after exposure to an audiovisual stimulus configuration where ITDs, but not ILS, were spatially unreliable (intercept of fit in Figure 6A, −0.33°, t = −0.92, *p* = 0.38; slope, 1°/°, t = 4.96, *p* < 0.001).

**Figure 6.**
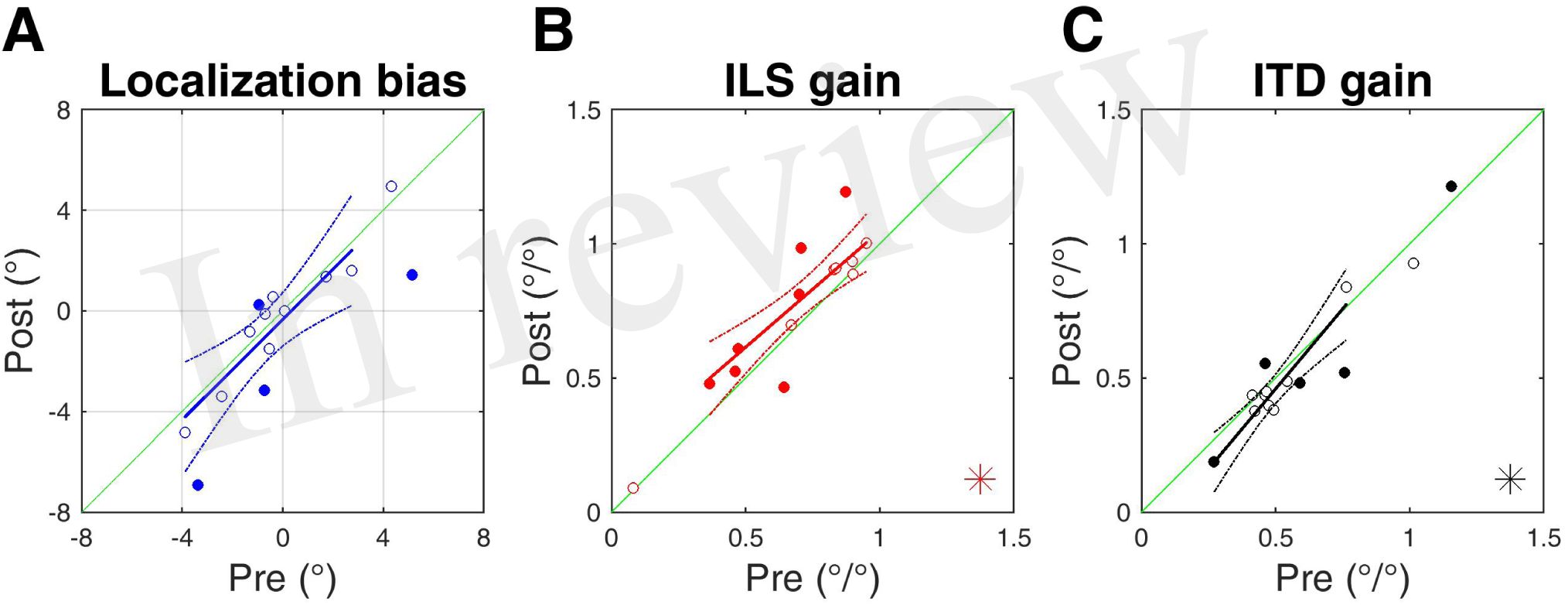
Subjects’ post-training multiple regression coefficients as a function of their pre-training regression coefficients for the random-ITD experiment. **(A)** Localization bias. **(B)** ILS cue gains. **(C)** ITD cue gains. Asterisks in the bottom right corner indicate significant group effects of training according to a linear mixed effects model. Solid symbols indicate significant within-subject changes (see Results). Solid lines are regressions of post-training score on pre-training score after outliers were excluded using a leverage test. Dashed lines are 95% confidence intervals for the fits.

Prior to audiovisual training, eight out of fourteen subjects exhibited higher spatial gains for ILS than ITDs, a difference that was significant (adjusted pre-test LME model estimates [and 95% confidence bounds]: ILS, 0.71°/° [0.58, 0.85]; ITD, 0.57°/° [0.45, 0.69]). As a result of pairing the ILS cue with a spatially congruent visual stimulus, twelve subjects showed ILS gain increases and in six subjects this change was significant (Figure 6B). The group increase was also significant (pre-training ILS gain, 0.67 ± 0.25°/°; post-training 0.75 ± 0.29°/°; t(3494) = 6.14, *p* < 0.0001). The confidence bounds [0.6, 1.13] for the slope of the regression (0.86) shown in Figure 6B encompass 1, indicating that the magnitude of the increase in spatial gain was constant, regardless of the initial value. Following audiovisual training, the ITD spatial gains decreased for ten subjects (three significantly) and increased in four subjects (two significantly) (Figure 6C). We observed a significant overall mean reduction in ITD gain between pre-(0.59 ± 0.25°/°) and post-training values (0.55 ± 0.27°/°; t(3494) = −3.58, *p* < 0.001), although again the slope of the regression shown in Figure 6C (1.19, [0.85, 1.54]) is not significantly different from 1, implying that the magnitude of the reduction was not related to subjects’ starting ITD gains.

### Audiovisual training with congruent visual and ITD cues and randomized ILS (random-ILS)

We next performed the complementary manipulation in cue reliability to the previous experiment by matching the visual stimulus position with the ITD cues, and randomizing the ILS position for each stimulus presentation. This was intended to increase the reliability of ITDs, whilst degrading the reliability of ILS. We hypothesized that this would result in analogous changes to those predicted for the random-ITD experiment, with any corrective changes in initial localization biases being accompanied by an increase in ITD gain and a decrease in ILS gain, producing an increase in the relative weighting of the more reliable ITD cue.

Eight out of fourteen subjects experienced significant changes in their localization biases in response to this training paradigm, and for six of those the absolute bias was lower after audiovisual exposure (Figure 7A). Although the mean group bias shift was not significant (pre-training bias, 0.17 ± 2.15°, post-training bias, −0.45 ± 2.36°; LME: t(3494) = - 0.16, *p* = 0.41), with outliers removed the regression shown in Figure 7A has a significant negative intercept (i.e., indicates a leftward shift in bias on average: intercept = −1.15, [−2.11, −0.19]; t = −2.77, *p* < 0.05). The slope of this regression is not significant (0.06°/°; t = 0.22, *p* = 0.83), which might indicate, as in the AVCON experiment, that the largest bias changes were seen in those subjects who already had the largest biases before training. This notion is supported by the fact that there is a negative correlation between subjects’ initial global localization biases and the direction and magnitude of their bias shifts (r = −0.79, *p* < 0.01; see also Figure 9A), implying that despite the overall leftward shift, the subjects who were most biased before audiovisual training showed the largest bias changes. However, the root mean square error (RMSE) of the fit shown in Figure 7A is almost double that shown in Figure 5A, indicating that randomizing the position of the ILS cue led to localization bias changes that were much more variable compared with the experiment where vision matched the position of both binaural cues (AVCON; reduction in localization bias).

**Figure 7.**
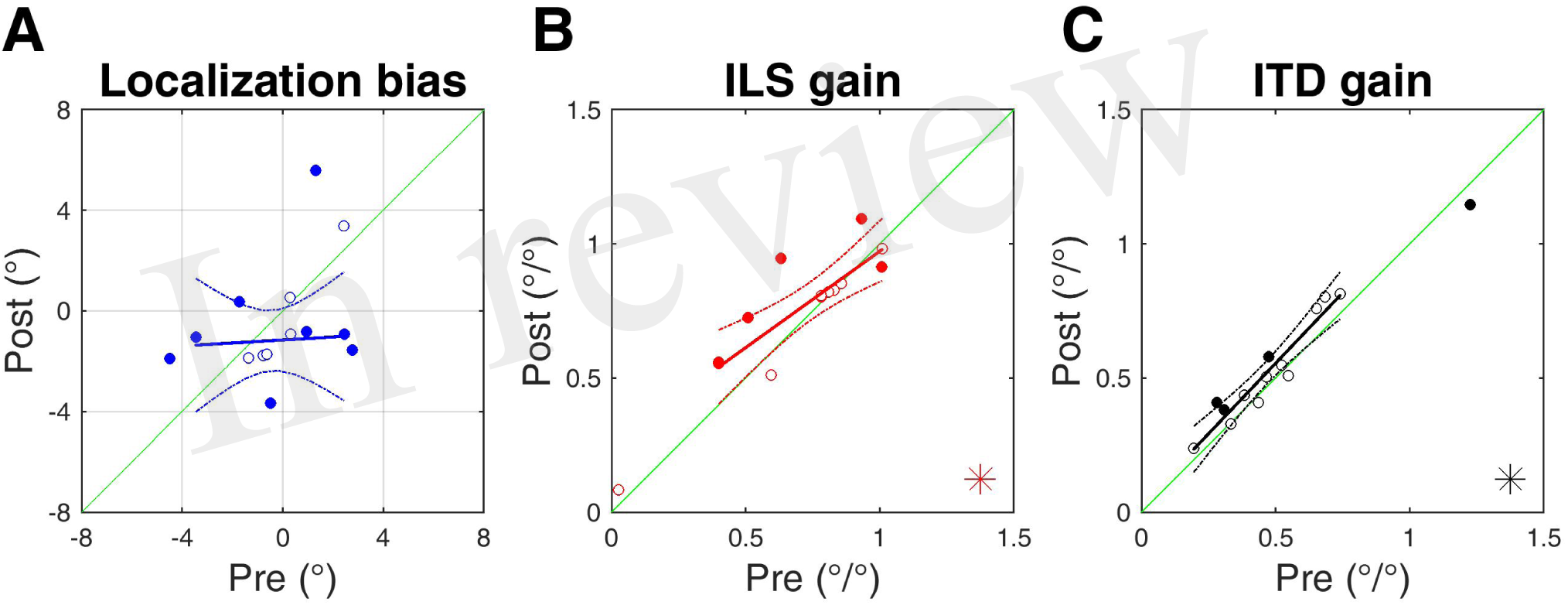
Subjects’ post-training multiple regression coefficients as a function of their pre-training regression coefficients for the random-ILS experiment. **(A)** Localization bias. **(B)** ILS cue gains. **(C)** ITD cue gains. Asterisks in the bottom right corner indicate significant group effects of training according to a linear mixed effects model. Solid symbols indicate significant within-subject changes (see Results). Solid lines are regressions of post-training score on pre-training score after outliers were excluded using a leverage test. Dashed lines are 95% confidence intervals for the fits.

As before, pre-training spatial gains were higher for ILS than ITDs in most subjects (n = 11/14), and as a group the difference was significant (adjusted pre-test LME model estimates [and 95% confidence bounds]: ILS, 0.72°/° [0.59, 0.86]; ITD, 0.53°/° [0.41, 0.66]). Surprisingly, nine out of fourteen subjects increased their ILS gains (five significant; Figure 7B), and a group increase in gain was observed (pre-training gain, 0.68 ± 0.28°/°; post-training, 0.75 ± 0.25°/°; t(3494) = 4.01, *p* < 0.0001). However, the confidence bounds for the regression slope shown in Figure 7B show that it is significantly lower than 1 (0.72°/°, [0.45, 0.99]), suggesting that subjects with the lowest starting gains exhibited the largest changes while those with high starting gains changed little.

Ten subjects increased their ITD gain after audiovisual training, although the change was only individually significant in three cases (see Figure 7C). The group increase in ITD gain was also significant (pre-training gain, 0.52 ± 0.26°/°; post-training, 0.56 ± 0.24°/°; t(3494) = 3.6, *p* < 0.001), but the confidence bounds for the slope of the regression shown in Figure 7C encompass 1 ([0.84, 1.26]), indicating that the magnitude of ITD gain increase was constant, regardless of subjects’ starting gains. There was no correlation between the change in localization bias and the change in either ILS (r = −0.1, *p* = 0.74) or ITD (r = −0.003, *p* = 0.99) cue gains.

### Audiovisual training with spatially disparate auditory and visual cues (VAE)

In this experiment, all sound localization cues were offset from the visual stimulus by 10° to either the left or the right in order to induce the ventriloquism illusion. Every subject reported being unaware of the disparity when questioned after testing. We first standardized the side of the VAE by reversing the sign of ILS angles, ITD angles and responses for those subjects who had performed this task with the visual stimulus positioned to the right of the auditory stimulus. In Figure 8A, the presence of a VAE shift is therefore indicated by a shift leftwards in a subject’s multiple regression intercept (i.e., by data-points falling below the green x = y line following training). Apart from three subjects whose bias shifted away from the direction of the visual stimulus (two significant), all of the data-points are below the x = y line and in nine of these eleven subjects this shift was significant, as was the group mean change (pre-training bias, −0.14 ± 2.97°; post-training bias, −1.81 ± 2.7°; t(3494) = −8.42, *p* < 0.0001). When we examined only subjects who displayed a significant shift in the direction of the previously presented visual stimulus, the mean VAE size was 2.92 ± 0.93°. The regression slope shown in Figure 8A does not differ significantly from 1 (0.68, [0.2, 1.15]), indicating that, in general, the shift magnitude was independent of any initial localization bias.

**Figure.8.**
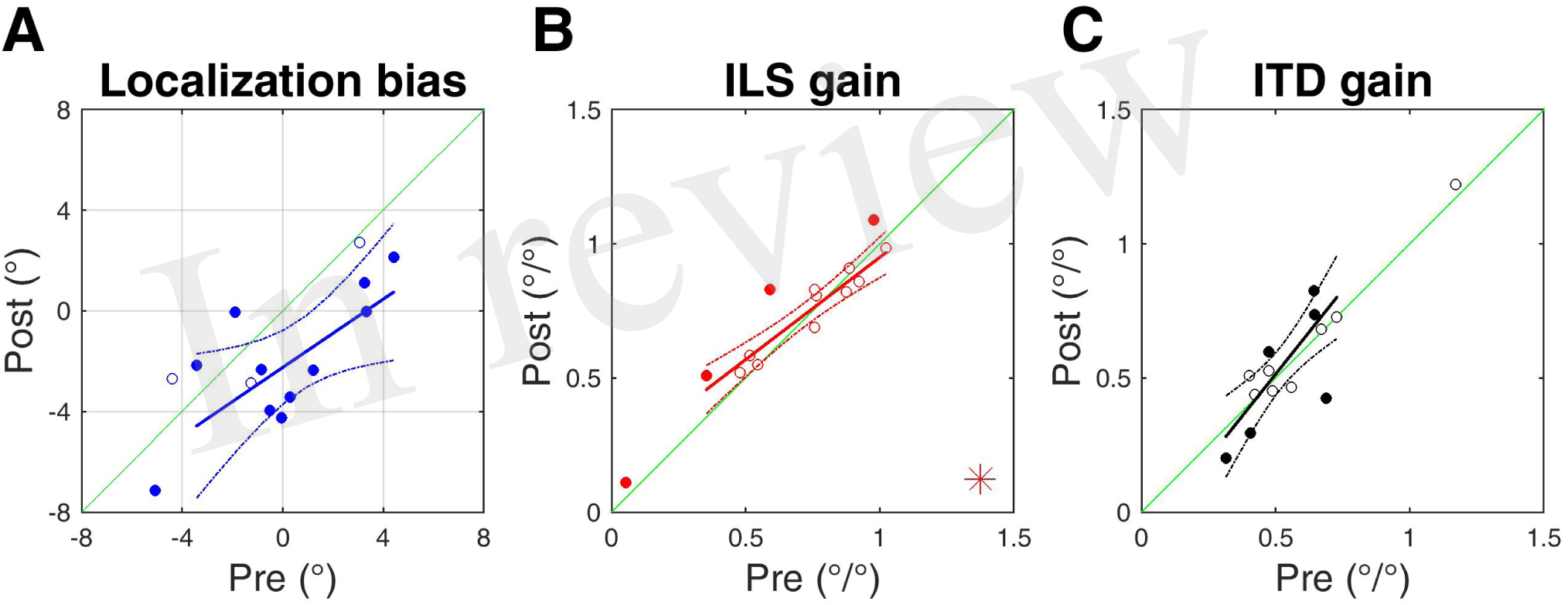
Subjects’ post-training multiple regression coefficients as a function of their pre-training regression coefficients for the VAE experiment. Stimulus values and responses have been reversed for subjects who experienced a visual stimulus that was offset to the right of the auditory stimulus during training. **(A)** Localization bias. **(B)** ILS cue gains. **(B)** ITD cue gains. Asterisks in the bottom right corner indicate significant group effects of training according to a linear mixed effects model. Solid symbols indicate significant within-subject changes (see Results). Solid lines are regressions of post-training score on pre-training score after outliers were excluded using a leverage test. Dashed lines are 95% confidence intervals for the fits.

Prior to audiovisual training, ten out of fourteen subjects had higher spatial gains for ILS than ITDs, although the confidence intervals for the ILS overlap the ITD estimate (adjusted pre-test LME model estimates [and 95% confidence bounds]: ILS, 0.7°/° [0.57, 0.83]; ITD, 0.58°/° [0.47, 0.69]). After training with spatially disparate auditory and visual cues, ten subjects showed an increase in their ILS gain, and four did so significantly (Figure 8B). The group effect (pre-training ILS gain, 0.68 ± 0.27°/°; post-training, 0.72 ± 0.25°/°) was also significant (t(3494) = 2.8, *p* < 0.01). However, the confidence bounds [0.6, 0.93] for the slope (0.76) of the regression shown in Figure 7B are less than 1, implying that, in this experiment, a larger gain increase was seen in subjects whose ILS gains were initially low, whereas those with higher initial ILS gains changed little or decreased their ILS gain.

Eight subjects increased their ITD gains and in three subjects this change was significant, although this was also true of three subjects whose ITD gains decreased. As a group (pre-training ITD gain, 0.58 ± 0.21°/°; post-training 0.58 ± 0.25°/°) there was no significant change (t(3494) = 0.6, *p* = 0.55) (Figure 8C). We observed no correlation between the change in localization bias, i.e., the magnitude of the VAE, and the change in either ILS (r = 0.41, *p* = 0.14) or ITD (r = −0.27, *p* = 0.35) cue gain, and four subjects who did not exhibit significant changes in the gains of either cue nonetheless displayed significant shifts in localization bias. Thus, visually-induced remapping of auditory space was not always accompanied by changes in cue gain.

### Changes in variability of sound localization bias

The experiments described above demonstrated that where one or both binaural cues were aligned with the visual stimulus (AVCON, random-ITD and random-ILS), only the AVCON and random-ILS configurations produced significant bias reductions in a substantial number of subjects (AVCON, n = 7; random-ILS, n = 6). Furthermore, the flat slopes for both fits in Figures 5A and 7A imply that, in both cases, larger bias reductions occurred for those subjects who were most biased to begin with. We confirmed this by removing outliers and fitting the regressions shown in Figure 9A. Although the slopes are similar for the two significant fits, the Pearson’s correlation coefficient for the random-ILS configuration is lower than for AVCON, reflecting the higher pre-training/post-training regression RMSE for random-ILS (Figure 7A) compared with AVCON (Figure 5A). We further examined the group bias scores for reductions in variability between pre-test and post-test by subjecting them to paired variance ratio tests. The variances of the pre- and post-test localization biases for each experiment are shown in Figure 9B. A significant reduction in variance was observed only in the AVCON configuration, where vision was spatially congruent with both ILS and ITDs, according to both a parametric (Pitman-Morgan, pre-post variance ratio = 3.02, *p* < 0.05) and nonparametric (Grambsch, z = 2.15, *p* < 0.05) paired variance ratio test. This suggests that auditory spatial cues need to provide consistent spatial information for exposure to spatially-congruent visual stimuli to evoke improvements in sound localization at the group level. By contrast, while the VAE condition induced an adaptive shift in auditory bias, the variability of the subjects’ responses was unchanged.

**Figure 9.**
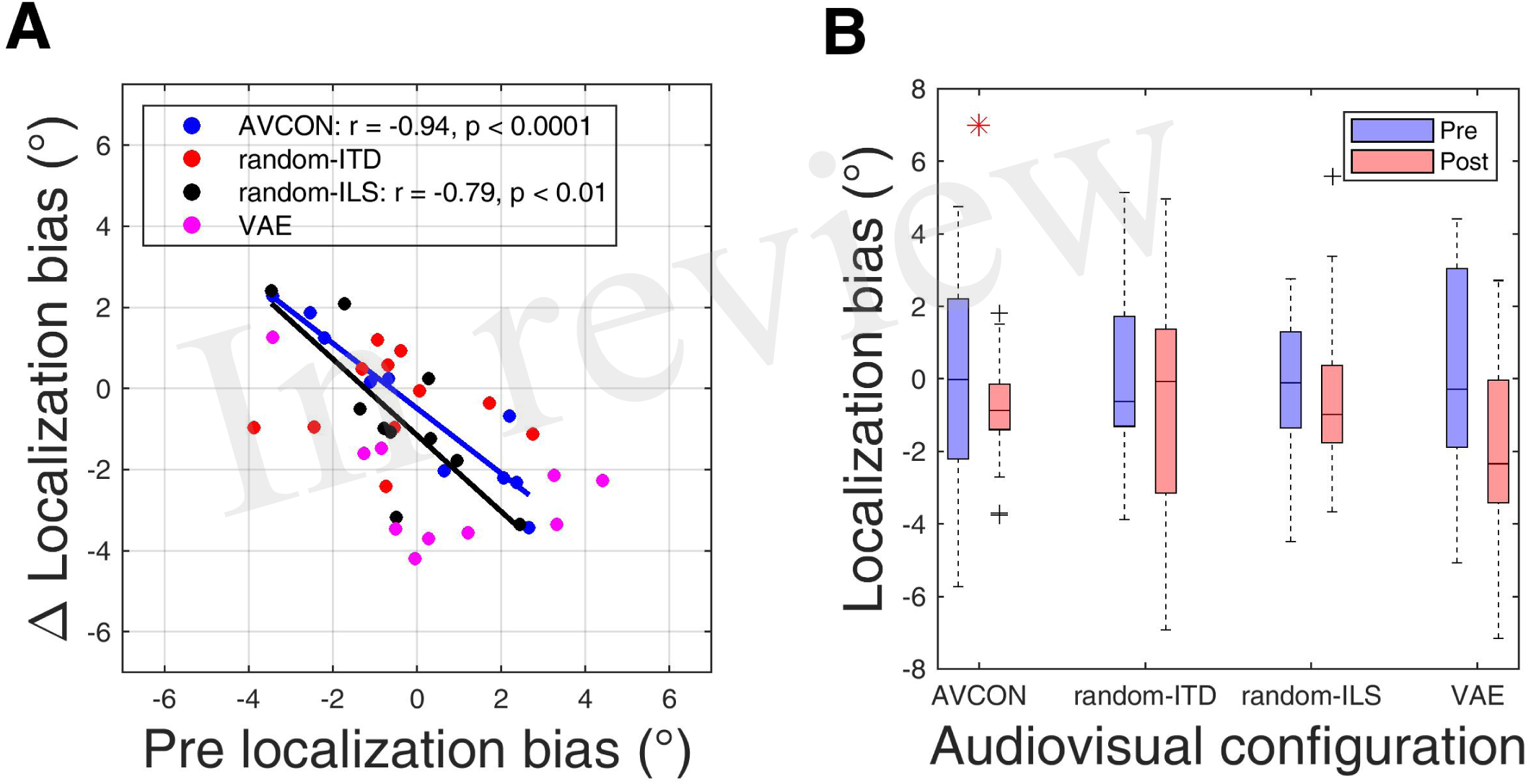
Changes in auditory localization bias. **(A)** Change in localization bias regressed on pre-training localization bias after outliers have been disregarded with a test of leverage. Regressions, Pearson’s *r* and their *p*- values are shown for experiments yielding a significant correlation between pre-training bias magnitude and the magnitude of the bias change induced by audiovisual training. **(B)** Distribution of all subjects’ localization biases before and after training. Significant reductions in sample variance induced by training are indicated with a red asterisk at the top of the panel.

### Changes in spatial gains of binaural cues

Figure 10 shows the absolute changes in ILS and ITD gains for each subject in each experiment, as well as the bootstrapped center of a 95% confidence ellipse around the data to illustrate the underlying pattern of spatial cue gain changes. When all auditory and visual cues came from the same position during audiovisual training (AVCON), subjects generally increased their ILS gain (maximum histogram bin centred at +0.12°/°), whilst maintaining their ITD gain at a constant value (maximum at +0.003°/°) (Figure 10A). Thus, spatial gains increased for the more salient ILS, but did not change for the less salient ITDs.

**Figure 10.**
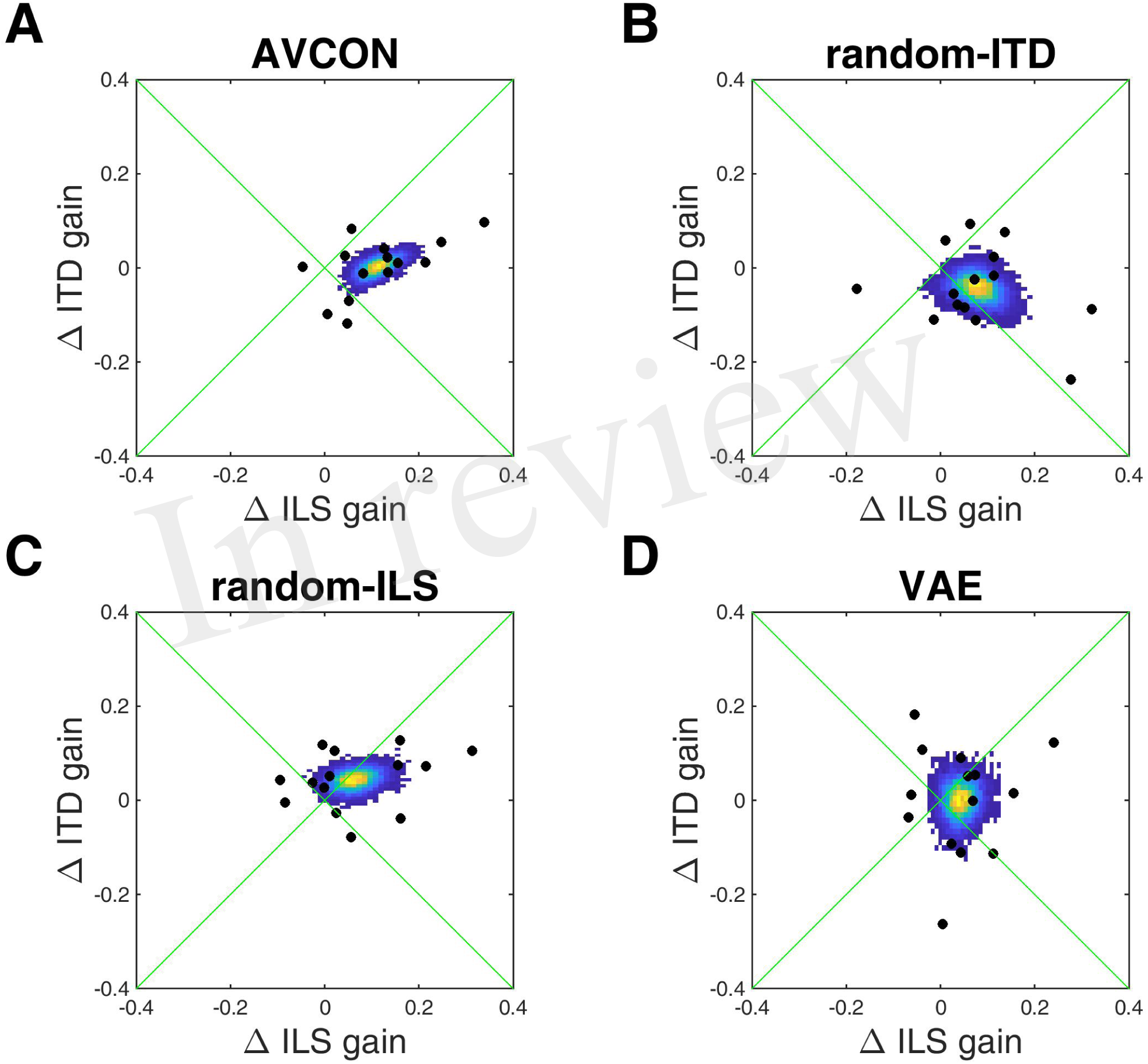
ILS and ITD cue spatial gain changes. The colored region indicates the bootstrapped origin of a 95% confidence ellipse fitted to the data; the yellow region indicates the most commonly observed values.

Matching the ILS cue with the visual cue whilst randomizing the ITD position (random-ITD) during training again resulted in an increase in gain for the reliable ILS cue (maximum bin value +0.08°/°), but this was now accompanied by a less pronounced decrease in gain for the unreliable ITD cue (ITD maximum bin value −0.05°/°) (Figure 10B). However, in contrast to the AVCON condition, where both auditory cues matched the location of the visual stimuli, we did not see an accompanying correction of subjects’ initial sound localization biases.

In the complementary audiovisual configuration where ITDs corresponded with visual cues and ILS were randomized (Figure 10C), comparable increases in gain were observed for both the ITD (+0.07°/°) and the ILS (maximum bin value +0.04°/°) cues. Thus, as in the random-ITD experiment, the spatial gain of the auditory localization cue (ITD in this case) that matches the position of the concurrent visual stimulus increases, but this time we also observed an increase in spatial gain for the randomized cue (ILS).

Finally, exposing subjects to consistent but spatially incongruent audiovisual stimulation (VAE) during training induced a broad range of gain adjustments (Figure 10D). Nonetheless, ILS spatial gains tended to increase (maximum bin value +0.04°/°), whilst the ITD gains remained relatively static (maximum +0.005°/°).

Taken together, these analyses suggest that the spatial gains of different binaural cues can be adjusted independently, dynamically and flexibly to reflect the salience and/or spatial reliability of those cues in the environment. However, if they are initially low, auditory cue gains can also undergo increases in the presence of spatially incongruent visual information.

### Changes in relative weighting of binaural cues

To understand how these changes in ILS and ITD spatial gains reflected changes in the relative weighting of each cue, we calculated a “binaural weighting” index for each subject (see Methods), for which values of −0.5 and 0.5 represent the extreme situations where either ITD or ILS dominate, respectively, and a value of 0 represents equal weighting between the cues. All four experiments yielded positive binaural weightings before training (pre mean ± SD: AVCON, 0.04 ± 0.17; random-ITD 0.03 ± 0.17; random-ILS, 0.07 ± 0.19; VAE, 0.04 ± 0.17), confirming that, as indicated by the spatial gain relationship between the cues, the ILS was generally the dominant localization cue. Paired t-tests revealed significant positive shifts in binaural weighting in the AVCON (post mean ± SD: 0.08 ± 0.16; t(13) = −5.72, *p* < 0.0001) and random-ITD (0.08 ± 0.18; t(13) = −3.32, *p* < 0.01) experiments, indicating that the dependence on the ILS cue was further increased by congruent visual stimulation. However, no net change in binaural weighting for the random-ILS (0.07 ± 0.16; t(13) = −0.29, *p* = 0.77) or VAE (0.06 ± 0.17, t(13) = −1.58 *p* = 0.14) experiments was observed.

More interesting are the relationships between subjects’ pre- and post-training binaural weights (after outlier removal), shown in Figure 11 for each audiovisual configuration. In the two experiments where the binaural weighting shifted significantly towards the already higher-weighted ILS cue (AVCON and random-ITD), a comparable shift was observed irrespective of subjects’ binaural weighting before training (AVCON, regression slope [and confidence bounds]: 0.94 [0.83, 1.05], Figure 11A; random-ITD: 1.03 [0.84, 1.22], Figure 11B). Furthermore, the magnitude of the weighting shift was similar for the two experiments (AVCON, intercept = 0.04 [0.03, 0.06]; random-ITD, 0.05 [0.02, 0.07]). Thus, in contrast to the spatial gain adjustments in these two experiments, the way the binaural cue weights changed was independent of contextual stimulus factors.

**Figure 11.**
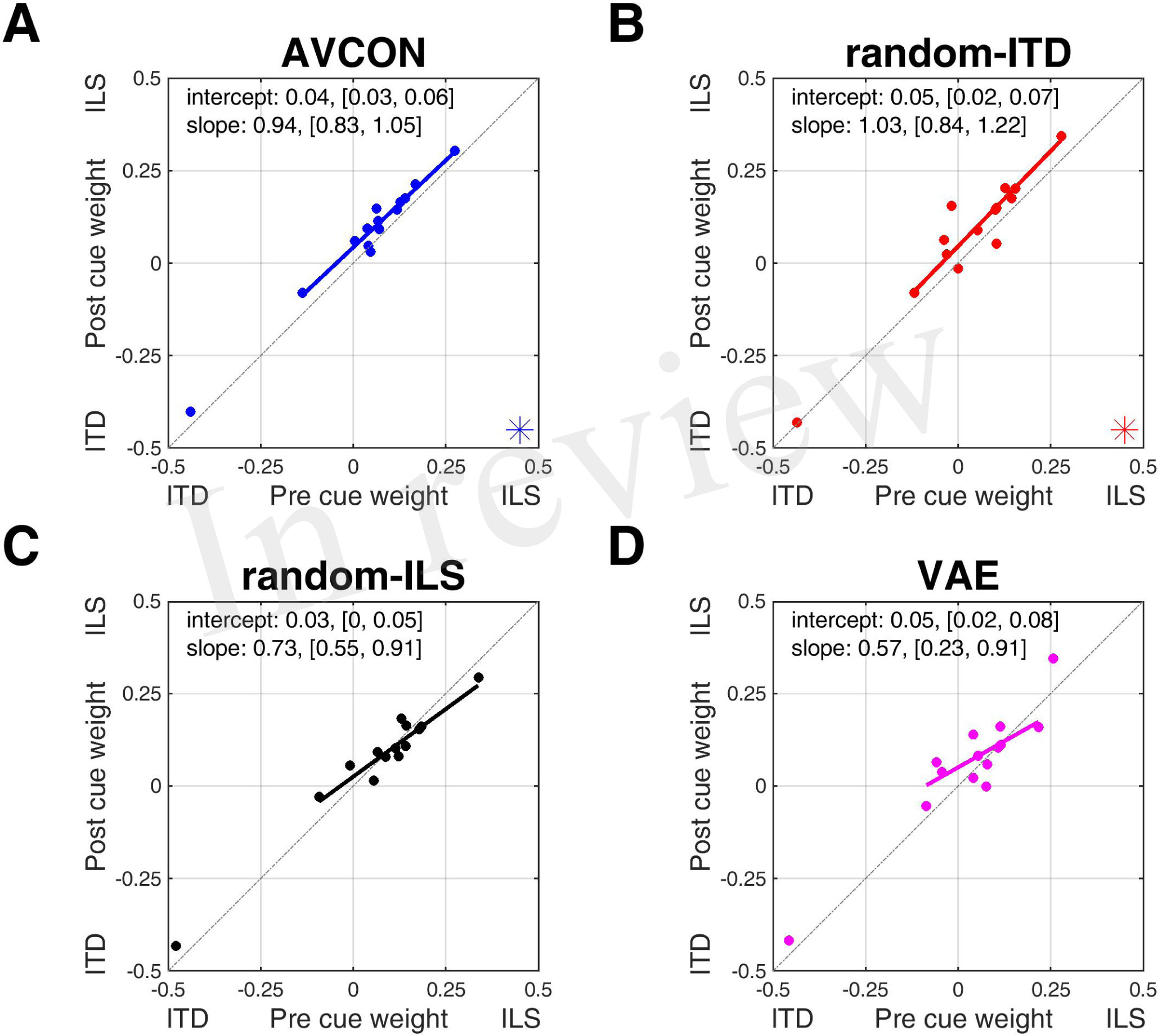
Binaural cue weighting changes for the four audiovisual configurations used in this study. Asterisks in the bottom right corner indicate significant group changes in binaural weight after training (see Results).

For the two experiments where the mean shift in binaural weighting was not significant (random-ILS and VAE), the slopes of the regressions of post-on pre-training binaural weights were significantly less than 1 (random-ILS, 0.73 [0.55, 0.91]; VAE, 0.57 [0.23, 0.91]), and they intersected the x = y line (i.e., no change was predicted between pre- and post-training scores) at similar binaural weighting values that slightly favored the ILS cue (random-ILS, +0.09; VAE, +0.12) (Figures 11C and 11D). It is possible that this effect results from the statistical phenomenon of regression to the mean (Bonate, 2000), although since we excluded outliers, and also did not observe similarly flattened slopes for the experiments where binaural weighting did change significantly, this seems unlikely. The data therefore suggest that for both of these audiovisual configurations, the direction of cue re-weighting depended upon whether subjects already weighted the more reliable ILS above or below a critical threshold before training: the higher the pre-training scores were above this value, the stronger the tendency for binaural weights to shift towards the ITD cue after training. Conversely, the lower the pre-training weights were below this threshold, the larger the binaural weighting shift in the direction of the ILS cue.

## DISCUSSION

In this study, we measured how human listeners weight different auditory spatial cues when they localize broadband sounds presented using non-individualized HRTFs. We manipulated the relative reliability of ITDs and ILS by pairing the sounds with visual stimuli under different conditions of audiovisual spatial congruence over the course of one hour. Our data show that a visual “teacher” signal induces corrective changes in auditory localization (characterized by individual reductions in bias and more uniform biases across subjects) only when it is spatially congruent with all available sound localization cues. In contrast, if the position of one auditory localization cue was randomized, consistent improvements in sound localization were not observed. Furthermore, in line with other studies of remapping of auditory space by vision (Radeau and Bertelson, 1974; Recanzone, 1998), we found that presenting auditory localization cues that were offset from a visual stimulus (VAE) induced a ventriloquism aftereffect of around 2°, corresponding to about 20% of the previous audiovisual spatial conflict, but no changes in the variability in bias across subjects.

These shifts in localization bias were accompanied by context-dependent changes in the spatial gains of the auditory cues. The gains of the initially more heavily weighted cue (in this case, ILS) increased when reinforced by spatially congruent visual signals (AVCON; random-ITD), but the gains of the initially less weighted cue (ITD) only decreased if the statistical unreliability of that cue was explicitly emphasized by randomizing its position relative to the visual stimulus (random-ITD). Nevertheless, the relative increase in ILS cue weighting in these two configurations was similar, indicating that unlike the spatial cue gains, the capacity for re-weighting may be limited and insensitive to contextual stimulus reliability. In the two experiments where visual cues and ILS provided conflicting spatial information (random-ILS, VAE), we only observed spatial gain increases for this cue if subjects’ pre-training gains were already low. That such a range of rapid changes can be achieved even when using another person’s ears illustrates the remarkable plasticity of the auditory system and has implications for auditory training schemes in clinical settings, as well as for achieving optimal human interaction with auditory displays in real and virtual environments.

### Audiovisual training reduces sound localization bias

We observed significant reductions in auditory spatial bias in most subjects, as well as in the range of bias values across subjects, when listeners carried out the oddball task with spatially congruent auditory and visual stimuli in the AVCON experiment. Because the virtual auditory space stimuli were presented using non-individualized HRTFs, we would expect the listeners’ localization judgments to be less accurate than if the stimuli had been based on measurements from their own ears (Wenzel et al., 1993; Middlebrooks, 1999; Mrsic-Flogel et al., 2001). Our results confirm findings from previous work showing that training with spatio-temporally congruent visual and auditory cues in listeners with abnormal or non-individualized auditory inputs produces improvements in sound localization (Isaiah et al, 2014; Trapeau and Schönwiesner, 2015; Stitt et al, 2019), and that these improvements can occur following limited exposure to multisensory inputs (Berger et al., 2018).

Exposure to spatially-congruent auditory and visual stimuli (AVCON) produced the largest corrective bias reductions in subjects with larger initial localization biases (Figure 9A), although there was some indication that they corrected to a point just to the left of the midline rather than at the midline itself. This may find an explanation in recent studies of pseudoneglect that demonstrate opposite spatial attentional biases for visual and auditory stimuli in healthy listeners, such that performance in spatial attention tasks is characterized by a leftward bias for visual stimuli (Loftus and Nicholls, 2012; Thomas et al., 2014, 2017) and a rightward bias for stimuli presented to the auditory modality (Sosa et al., 2010, 2011). These biases have been postulated to result from hemispheric asymmetries in controlling the deployment of spatial attention. Reports of a greater potency of visual capture of auditory cues in the left versus the right visual hemifield (Sosa et al., 2011) are also consistent with the tendency of several of our subjects to show a small leftward bias in sound localization following training with spatially congruent audiovisual cues.

### Changes in spatial gain and cue weighting

Our subjects generally weighted ILS higher than ITDs, which is consistent with a previous observation that the former may be more salient when binaural cues are presented over headphones in multisensory experiments (Kumpik et al., 2014). This is likely to be because our stimuli were bandpass filtered from 0.5-16 kHz, which would have removed some of the fine-structure ITD information from the HRTF and thus made the ITD less salient than might otherwise be expected (Wightman and Kistler, 1992).

We found that audiovisual stimulation changed the relationship in the brain between individual sound localization cues and spatial location, depending on their contextual reliability. In the case of fully congruent audiovisual information (AVCON), there was a trend for the largest increases in spatial ILS gain to be seen in subjects who had the highest ILS gains prior to training. That the spatial gains for the ITD cue remained static in this experiment contrasts with the decrease in gain observed when ITD position was made explicitly unreliable relative to vision in the random-ITD experiment. In this case, the increases in ILS gain were reciprocated by decreases in ITD spatial gain (Figure 10B). Thus, in mapping physical sound localization cues to external spatial positions the brain dynamically takes account of the contextual reliability of all the available information (Rohe and Noppeney, 2015) and adjusts the spatial gains of cues accordingly. These two configurations also led to significant up-weighting of ILS over ITDs, but since the magnitude of the change was the same in both experiments, it is possible that the extent to which binaural cue weighting can occur may be limited.

That the relative weighting of auditory localization cues can be changed by experience is consistent with previous work demonstrating that spatial hearing can adapt to altered auditory inputs by up-weighting whichever cues are deemed to be the most reliable for localization and down-weighting cues that provide conflicting or unreliable information (Kacelnik et al, 2006; van Wanrooij and van Opstal, 2007; Kumpik et al, 2010; Keating et al, 2013, 2015, 2016). Together, these findings suggest that the brain can re-weight auditory spatial cues in different ways depending on their relative reliability. We believe this to be the first direct demonstration of binaural cue reweighting in response to audiovisual training.

Surprisingly, the VAE and random-ILS experiments also induced significant group increases in ILS gain even though that cue was spatially incongruent with vision. Indeed, subjects with initially low ILS gains underwent the largest increases, which were unrelated to the observed bias changes, indicating that the degree of correspondence with the more reliable visual signal is not the only factor that drives plasticity in the neural processing of auditory localization cues. In the random-ILS experiment at least, the increase in ILS spatial gain may reflect the fact that subjects found this cue to be highly salient and difficult to ignore; for at least the first 2-3 oddball training blocks, they made involuntary eye movements toward the randomized position of the ILS cue at each stimulus presentation, likely reflecting sound-driven orienting responses that help to select relevant stimuli for spatial attention amongst competing distractors (Krauzlis et al., 2013). However, neither of these experiments induced significant re-weighting of ILS and ITDs, again indicating that whilst spatial cue gains show considerable flexibility in their plasticity, this does not necessarily translate to changes in how the cues are weighted by the brain.

### Auditory bias shifts and cue re-weighting are fast and independent processes

Our data show that remapping of auditory space (i.e., shifts in localization bias) and re-weighting of auditory localization cues are reasonably rapid, with both changes being apparent after an hour of audiovisual training. Although it has been shown that remapping of auditory space can manifest after only milliseconds of exposure to an audiovisual disparity (Wozny and Shams, 2011b), it remains to be seen whether a similar time-scale applies to the re-weighting of binaural cues. At the group level, we observed localization bias shifts both with and without cue re-weighting, and in the random-ITD configuration cue re-weighting occurred in the absence of any shifts in localization bias. While changes in auditory localization bias can be interpreted in terms of a reduction in audiovisual spatial conflict, the weights given to individual sound localization cues may change in different ways depending on their saliency and contextual reliability, with the strongest positive cue re-weighting effects observed for reliable localization cues containing a rich spectral profile. That re-weighting and remapping of different sound localization cues can take place independently has also been demonstrated in experiments in which human listeners are trained to adapt to a temporary unilateral hearing loss (Keating et al., 2016).

### Neurophysiological basis for plasticity

Studies in patients with brain lesions suggest that short-term auditory adaptation to spatially congruent and incongruent audiovisual stimuli is subserved by distinct neural circuits (Passamonti et al., 2009; Bertini et al., 2016), with congruent stimuli activating a circuit involving the superior colliculus (SC) and extrastriate visual cortex, resulting in a reduction in position-specific auditory localization error. Conversely, the geniculostriate pathway has been implicated in a generalized recalibration of auditory space for incongruent stimuli, probably as a result of a direct biasing of auditory cortical activity by visual signals (Bonath et al., 2007, 2014; Bertini et al., 2010).

Both the re-weighting of spectral cues (Keating et al., 2013) and adaptive shifts in sensitivity to ILDs (Keating et al., 2015) take place in the primary auditory cortex of ferrets raised with one ear occluded, with largely separate populations of neurons undergoing each form of plasticity (Keating et al., 2016). Similarly, adaptation to modified ITDs in adult humans is associated with changes in auditory cortical activity (Trapeau and Schonweiser, 2015). Under normal hearing conditions, spatial processing in the auditory cortex is enhanced by congruent visual inputs (Bizley and King, 2008) and by performance of a visual task requiring subjects to direct their attention to the left or right (Salminen et al., 2013). While consistent with the possibility that visual updating of the perceived direction of a sound source is based on changes in auditory cortical activity, it is also possible that interactions between visual and auditory inputs in the midbrain may be involved. Reciprocal connections exist between the SC and different regions of the inferior colliculus (IC) (Doubell et al., 2000; Stitt et al., 2015), providing a source of retinotopically organized input into the auditory midbrain. Furthermore, auditory responses in the monkey IC are modulated by changes in gaze direction (Zwiers et al., 2004; Gruters and Groh, 2012), implying a role for the midbrain in coordinating neural representations of auditory and visual space. In this regard, it is interesting to note that IC neurons can register when auditory localization cues become misaligned, with neurons in the nucleus of the brachium of the IC, which projects topographically to the SC (King et al., 1998), being tuned to natural combinations of these cues (Slee et al., 2014). Together with evidence that auditory corticocollicular neurons play a critical role in training-dependent reweighting of spatial cues in monaurally-deprived adult ferrets (Bajo et al., 2010), these findings suggest an additional midbrain site for the visual calibration of auditory space.

In order to evaluate a cue’s reliability for re-weighting, a history of its spatial congruence with concurrent visual information must be established. Given evidence that ventriloquism can act at early, preattentive processing stages (e.g., Passamonti et al., 2009; Bertelson and Aschersleben, 1998; Vroomen et al., 2001) and that ventriloquism aftereffects can occur after a single trial (Wozny and Shams, 2011b), it is likely that automatic online capture of auditory cortical population responses by visual inputs affects subsequent spatial processing depending on the statistics of audiovisual congruency from both the immediate and cumulative past (Bruns and Röder, 2015). Thus, even in the AVCON experiment, subjects probably underwent ventriloquism and its aftereffects for a period of time before their auditory responses became aligned with the visual stimulus. How cortical and midbrain circuits interact to bring about visually-induced shifts in perceived sound-source location and changes in the underlying auditory localization cue weights remains to be determined.

In summary, our results show that audiovisual training can induce a recalibration of auditory space, which is associated with changes in the weighting of the cues that provide the basis for sound localization. Reliable auditory cues that are reinforced by spatiotemporally congruent visual stimuli have their weights increased, whereas cues that lack a consistent relationship with the visual stimuli may have their weights reduced, particularly if they are less salient to begin with. This highlights a potentially useful therapeutic avenue for using audiovisual training to refine the sound localization abilities of hearing-impaired listeners who are deprived of access to certain cues as a result of the frequency dependence of their hearing loss or the limitations of the devices used to treat them.

## Acknowledgments

We are grateful to Nicola Gillen for assistance with the data collection and thank our test subjects for their participation. Research funded by a Wellcome Principal Research Fellowship (WT108369/Z/2015/Z) and by the National Institute for Health Research (NIHR) Oxford Biomedical Research Centre (BRC). The views expressed are those of the author(s) and not necessarily those of the NHS, the NIHR or the Department of Health.

## Author Contributions

DK, JS and AK conceived and designed the study; DK and CC carried out the experiments and analyzed the data; DK, JS and AK wrote the manuscript.

